# SEQUIN: rapid and reproducible analysis of RNA-seq data in R/Shiny

**DOI:** 10.1101/2022.02.23.481646

**Authors:** Claire Weber, Marissa B. Hirst, Ben Ernest, Hannah Baskir, Carlos A. Tristan, Pei-Hsuan Chu, Ilyas Singeç

## Abstract

SEQUIN is a new web application (app) that allows fast and intuitive RNA-sequencing data analysis for organisms, tissues, and single cells. Integrated app functions enable uploading datasets, quality control, gene set enrichment, data visualization, and differential gene expression analysis. We also present the iPSC Profiler, a practical tool that helps to measure pluripotency and cell differentiation. Freely available to the public, SEQUIN empowers scientists to investigate transcriptome data firsthand with cutting edge statistical methods.

## Main

RNA-sequencing (RNA-seq) has become the method of choice for gene expression profiling and single-cell analysis (scRNA-seq)^1^. Massively parallel high-throughput sequencing is increasingly accessible and affordable. However, as new sequencing technologies and statistical methods rapidly evolve^2–4^, each new approach requires learning a new programming language or software tool. Another challenge is the need to merge independent sequencing libraries, which requires batch correction^5^. Here, we developed SEQUIN guided by the NIH principles of scientific data management (findability, accessibility, interoperability, reusability)^6^. SEQUIN is a R/Shiny app for realtime analysis and visualization of bulk and scRNA-seq raw count and metadata (**Fig. 1a**). The app was developed for novice users enabling complex data analysis, interactive exploration, and generation of publication-ready figures in one place. SEQUIN is open source and currently the most comprehensive platform for web browser-based gene expression analysis (https://sequin.ncats.io/app/).

**Fig. 1:**
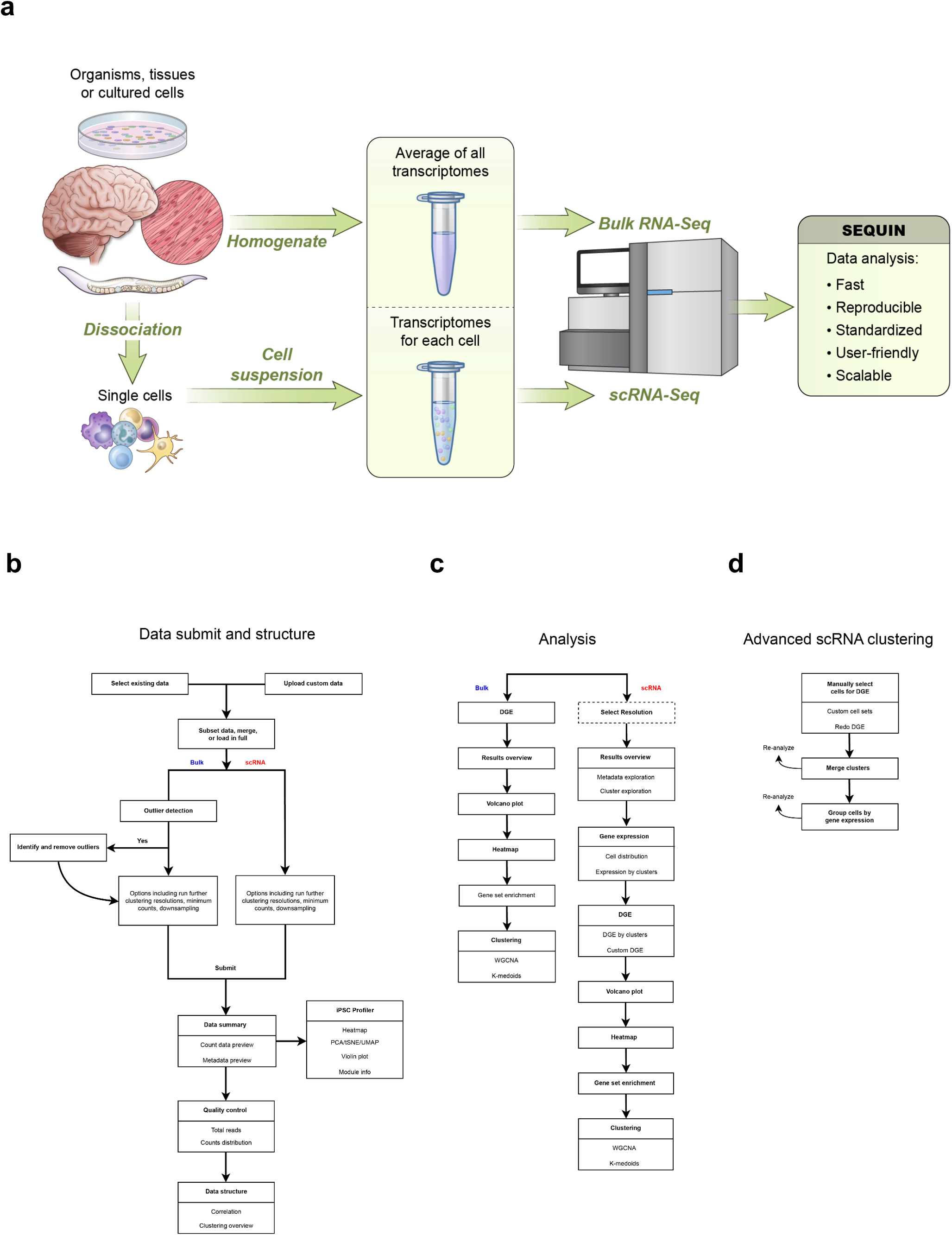
Versatility of SEQUIN for bulk and single-cell RNA-Seq. **a,** Overview of experimental models and next-generation sequencing that generate transcriptomic datasets. **b,** Workflow for the data submit and structure sections of the app. **c,** Workflow for analysis sections. The select resolution (in dashed box) is only available when the user selects “Run multiple resolutions using Seurat.” **d,** Workflow for advanced scRNA clustering with key features and options.

To illustrate the workflow, we first analyzed RNA-seq data from bulk samples (dataset ISB003_WA09) established from human embryonic stem cells (hESCs) and their progeny differentiated into ectoderm, mesoderm, and endoderm^7^. We uploaded the entire dataset and removed pseudogenes, ribosomal and mitochondrial genes, and genes with row sums less than 10 and transformed the data for normalization (log2(count +1)(**Fig. 1b**). The user can visualize a portion of the count matrix, the metadata, and visualize the distribution of samples. A total of 16 samples were analyzed (n = 4 for pluripotent and differentiated lineages) and 21,324 genes remained after filtering. The samples had relatively balanced counts per cell with an average of 21 million counts per sample (**Extended Data Fig. 1a**). Principal component analysis (PCA) revealed clustering of replicates by sample with 51% and 33% of the variance explained by PC1 and PC2, respectively (**Extended Data Fig. 1b**). We performed differential gene expression (DGE) analysis using pairwise two-group comparisons with DESeq2 with a minimum fold-change of 1 and an adjusted p-value of 0.05 (**Fig. 1c**). Of note, several other modeling options are available in the app including edgeR and limma-voom. These choices automatically generate a preview of the linear model design next to the “Submit” button (**Extended Data Fig. 2a**). Because no batch effect across these samples was observed, we did not batch correct (see Methods). The output for DGE is a table and plot of the total number of genes that are up- or down-regulated (**Extended Data Fig. 2b**). The volcano plot tab provides options to view volcano or MA plots as well as a significant filtered DGE table (**Extended Data Fig. 2c**). Guided by pairwise DGE comparisons, we explored upregulated differentially expressed genes (DE genes) and confirmed the expression of lineagespecific genes. Lineage-specific DE genes were significantly upregulated in pluripotent and differentiated cells. It is also possible to view a heatmap of the top 100 DE genes by contrast or by selecting a custom gene list. Cells can also be clustered by gene expression using two unsupervised approaches: weighted correlation network analysis (WGCNA)^8^ or k-medoids^9^ (**Fig. 1c** and **Extended Data Fig. 3**). These options are also available for scRNA-seq as described below.

To analyze scRNA-seq data, we used a recently published dataset (IS006_WA09) including pluripotent and differentiated cells^7^. Prior to data submission to the server, the full set of cells was randomly downsampled from 16,582 to 10,000 cells and all ribosomal RNA, mitochondrial, and pseudogenes were filtered out. The option to downsample was implemented to reduce count matrix load time and the RAM constraints in R. If the user prefers not to downsample and is willing to wait longer, the entire dataset can be uploaded (data is only stored for the duration of the app session). The standard Seurat workflow^10^ is incorporated into SEQUIN with options to adjust the number of PCA dimensions either by setting a cumulative percentage of variance in the PCs or using the default of 75% and the range of clustering resolutions (**Fig. 1b**). Prior to performing downstream analyses, the user can create a snapshot of the count matrix, the metadata, and the distribution of cells or samples after the dataset has been fully loaded. The samples had relatively balanced counts per cell (median 10,000) and the threshold for minimum counts removed poor quality or empty cells. Overall variance between samples can be visualized. Moreover, review of dimensionality reduction by selected metadata factor is possible with PCA, t-stochastic neighbor embedding (tSNE), and uniform manifold approximation projection (UMAP). Four distinct cell clusters were present in the tSNE plot (**Extended Data Fig. 4**).

We submitted the scRNA-seq dataset with varied resolutions from 0.1 to 1 by steps of 0.1; higher resolutions generate more clusters. Four to 16 clusters were created for resolutions 0.1 to 2, respectively. Based on the UMAP (**Fig. 2a**), Clustree plot (**Fig. 2b**), and silhouette plot (**Fig. 2c**) we concluded that 0.1 was the optimal resolution for clustering the different developmental lineages (**Fig. 1c**). Clustree is helpful to identify the best resolution for cell clustering based on the distribution^11^. Each resolution is represented by a row, with circle sizes corresponding to the total number of cells, and arrow thickness indicating the proportion of cells that flow from one cluster to another. Over-clustering occurs when there is frequent crossover of cells from one cluster to another at higher resolutions, and the user should choose a stable resolution lower than this point (**Fig. 2b**, rows 5-10). In the silhouette plot (**Fig. 2c**), the positive silhouette widths indicate that a given cell is closer to other cells within that cluster than to other clusters, which is ideal^12^. Together with the dimensionality reduction (**Fig. 2a**) and clustree plots (**Fig. 2b**), we confirmed that 0.1 resolution clearly separated cells. The UMAP revealed that four clusters separate cells by differentiation stage, which further supports the 0.1 resolution for clustering. The metadata can be explored in various other ways (**Fig. 1c and Extended Data Fig. 5a,b**).

**Fig. 2:**
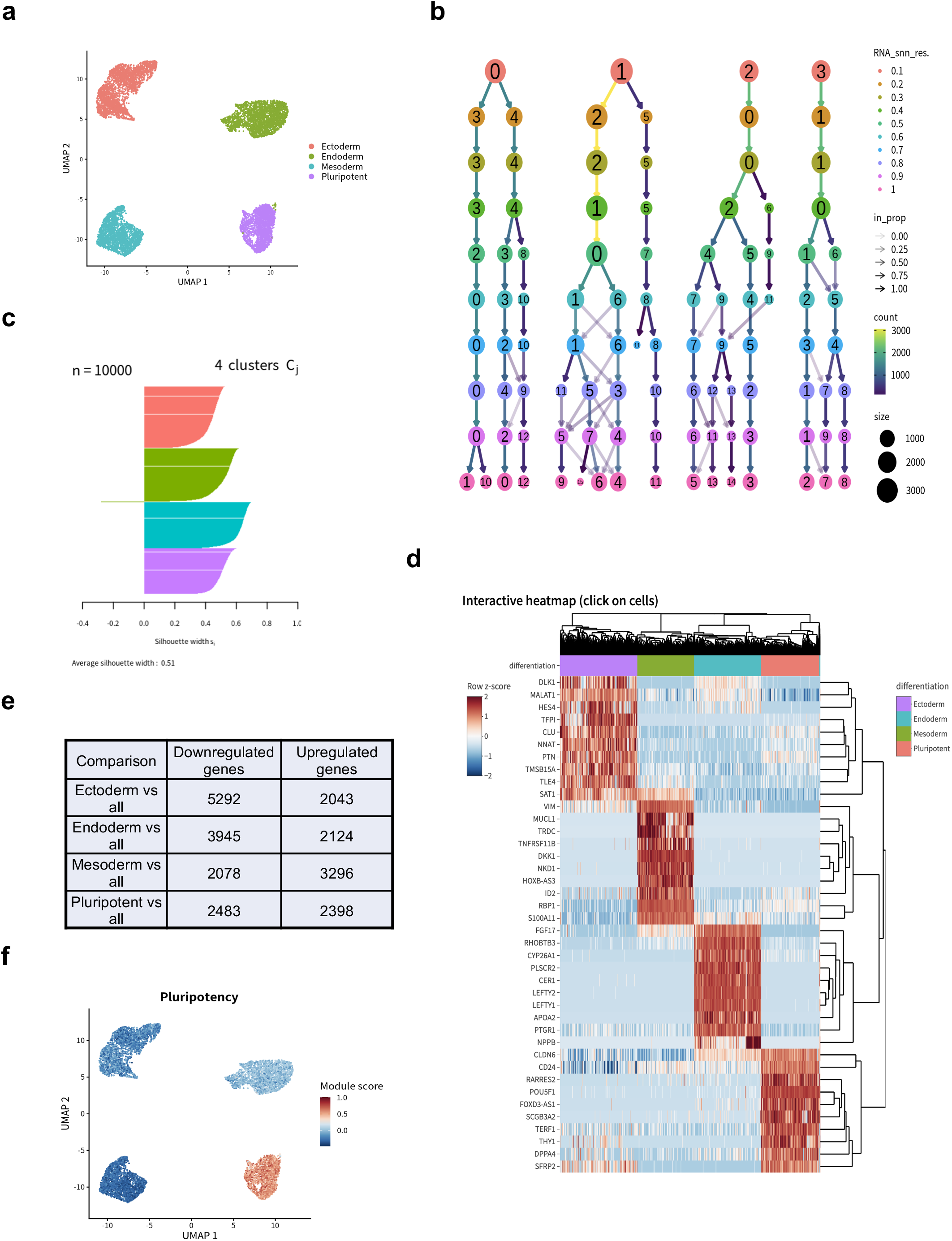
Analysis of an iPSC line to three primary differentiation lineages. **a**, UMAP clearly separates clusters by differentiation stage. **b**, Clustree flowchart identifies how cells sort into clusters at various resolutions. **c**, Silhouette plot cleanly separates clusters at 0.1 resolution. **d,** Interactive heatmap with a custom set of lineagespecific genes. **e**, Total up- and downregulated DE genes by differentiation stage as compared to the rest of the clusters. **f**, UMAP plot colored by pluripotency module scores reveals high scores in hESCs (WA09).

Prior to performing DGE, we inspected lineage-specific genes using the selection from the drop-down menu of available genes. Gene expression can be displayed by cluster or sample name in the form of bar plots and in PCA, UMAP, and tSNE (**Fig. 1c**). Example UMAPs are shown for POU5F1, HES4, DKK1, and PTGR1 (**Extended Data Fig. 6a-h**). SEQUIN also provides alternative options for DGE analysis (**Fig. 1c**). The user can compare a given cluster to the rest or select a factor in the metadata and the desired comparisons (**Fig. 1c**). Using an interactive UMAP, tSNE, or PCA plot, the user can manually lasso-select cells to perform DGE and gene set enrichment analysis (GSE) and download these tables (**Fig. 1d**). Lasso selection of cells to define new, custom clusters based on the user’s knowledge is a powerful feature of SEQUIN. To identify the uniquely DE genes per cluster, Custom DGE was performed (**Fig. 2e**) and lineage-specific results were consistent with a recent report^7^. Several thousand statistically significant genes were up- or downregulated for pluripotent and differentiated cells. The user can explore pairwise DE genes in a volcano plot or by generating a DE genes table based on log-fold change and adjusted p-value thresholds. While we provide core DE analysis, there are many options to further explore these genes. The user can generate a heatmap of the top 50 DE genes or by specifying a custom list of genes across samples (**Fig. 1c**). A selected metadata grouping factor for cells can be overlaid onto the heatmap samples. For instance, we targeted a set of genes involved in cell differentiation that showed scaled, normalized gene expression varied greatly by lineage (**Fig. 2d**). Subsequently, we performed GSE (**Fig. 1c**) on the statistically significant DE genes that were upregulated in each respective cluster (lineage) versus the rest of the clusters (the top 100 DE genes upregulated in the selected cluster). GSE calls the Enrichr API^13^ with a mirror of all gene set libraries available from that resource, including the popular KEGG, Gene Ontology, and ARCHS4^14^. To our knowledge, the full integration of DGE and GSE analyses in one R\Shiny app is unique to SEQUIN.

A valuable app feature is the ability to merge existing clusters or cluster cells based on the mean, median, or sum of gene(s) expression (**Fig. 1d and Extended Data Figs. 7,8**). A resolution of 0.1 created four distinct clusters, clearly separated by tissue lineage. Alternatively, a resolution of 0.3 was able to further separate the endoderm and ectoderm clusters to give a total of six clusters (**Extended Data Fig. 7a,b**). If after exploring the clustree and other plots, it becomes clear that a different clustering would be optimal, the user can merge clusters, which can then be named and given a comment to describe the merge (**Extended Data Fig. 7c,d**). By clicking “Save to database”, the updated metadata will be saved in the Relational Database Service (RDS) and can be reloaded with the count data for downstream analysis (**Extended Data Figs. 7d and 8**). Only metadata changes to existing experiments will be saved; custom uploaded data will not be stored in the RDS.

The iPSC Profiler is a special feature of SEQUIN and was developed to measure pluripotency and embryoid body differentiation and can complement PluriTest and ScoreCard, which were developed based on microarrays and quantitative PCR^15–17^. The iPSC Profiler is a gene module scoring tool for both bulk and scRNA based on the Seurat AddModuleScore function and classifies single cells or whole samples as pluripotent, ectodermal, mesodermal, or endodermal (**Fig. 1b**)^16^. Module scores differ from the simple average of a gene set in that they are directly comparable across set sizes and contents. We ran the Profiler for dataset IS006_WA09, then generated a UMAP of the pluripotency module (**Fig. 2f**) and a reference UMAP colored by expected cell lineage (**Fig. 2a**). Module scores were highest for endoderm and pluripotency gene sets (**Fig. 2f and Extended Data Fig. 9a**). Ectoderm and mesoderm cells scored relatively low for their gene sets, with only a subset of the cells having high scores for these two modules (**Extended Data Fig. 9b,c**). The housekeeping gene module scores were consistent across all lineages (**Extended Data Fig. 9d**). Module scores were well matched to the ScoreCard modules for pluripotency, ectoderm, mesoderm, and endoderm (**Extended Data Fig. 9e**). Single-cell and bulk RNA-Seq datasets yielded similar DE genes. Cell typespecific genes were significantly DE at each stage of differentiation in both datasets. While the bulk data does not identically mirror enriched ARCHS4 gene sets, we saw overall consistency across differentiation stages. With the ease and convenience of gene module scoring, the iPSC Profiler highlighted successful differentiation into the three primary tissue lineages. In summary, SEQUIN empowers users with different backgrounds to perform customizable analysis of bulk and single-cell RNA-seq in realtime and in one location. We believe SEQUIN sets a new standard for RNA-seq analysis within the R/Shiny environment.

## Supporting information

Ext Data Fig 1

Ext Data Fig 2

Ext Data Fig 3

Ext Data Fig 4

Ext Data Fig 5

Ext Data Fig 6

Ext Data Fig 7

Ext Data Fig 8

Ext Data Fig 9

## Acknowledgements

This study was supported by the NIH Common Fund (Regenerative Medicine Program) and in part by the NCATS DPI Intramural Research Program. We thank Rancho Biosciences and the ITRB and Informatics Group at NCATS. We acknowledge the contributions of all SCTL scientists for suggesting feature enhancements and testing SEQUIN. This project used the computational resources of the NIH High-Performance Computing (HPC) Biowulf cluster for R pipeline development (http://hpc.nih.gov).

## Extended Data Figure Legends

**Extended Data Fig. 1: Bulk RNA-seq analysis**

**a,** Quality control step shows total reads by differentiation stage, which are consistently on average 21 million total reads by differentiation stage. **b**, PCA plot of samples and replicates shows that PC1 accounts for 51% and PC2 33% of the variance in the data, respectively. Samples and replicates separate strongly by PC1 and PC2 clustering by differentiation. Example dataset ISB003 (WA09) was analyzed.

**Extended Data Fig. 2: Example analyses of DE genes**

**a**, DGE analysis options for the bulk RNA-Seq analysis of ISB003 (WA09) showing two-group comparisons by differentiation using DESeq2. The adjusted p-value cut-off is 0.05 with a minimum fold change of 1 and the linear model ~differentiation. **b**, Total DE genes by linear model showing the total number of genes up- and downregulated in the ectoderm vs. endoderm comparison. **c**, Volcano plot of total up- and downregulated DE genes for the ectoderm vs. endoderm comparison.

**Extended Data Fig. 3: WGCNA and k-medoid clustering**

**a**, **b**, WGCNA clustering. The WGCNA gene dendrogram for IS006 (WA09) is based on the hierarchical clustering of all genes. Colors below the row and column dendrograms are the dynamic tree cuts, which indicate the size (total number of cells per cluster). The WGCNA topological matrix plot is the correlation between pairs of genes and pairs of gene modules. A k-medoids consensus matrix heatmap based on all genes and samples from IS006 (WA09) (b). **c**, k-medoids heatmap reflects an estimate of the similarity between pairs of genes.

**Extended Data Fig. 4: tSNE plot by original identity**

A resolution of 0.1 reveals four distinct cell clusters with pluripotent, ectodermal, endodermal, mesodermal signatures.

**Extended Data Fig. 5: Overview of scRNA metadata analysis**

**a**, Exploration of cluster sizes and quality. **b**, Identification of relationships between factors in the metadata.

**Extended Data Fig. 6: UMAP dimensional reduction plots and boxplots of gene expression by cluster**

**a**, **b**, Pluripotent cells (cluster 2) express POU5F1 (OCT4). **c**, **d**, HES4 is strongly expressed by ectodermal cells (cluster 0). **e**, **f**, Mesodermal cells express DKK1 (cluster 3). **G**, **h**, Expression of PTGR1 by endodermal cells (cluster 1).

**Extended Data Fig. 7: Merging scRNA clusters**

**a**, UMAP of further clustering at 0.3 resolution reveals six clusters, splitting endoderm and ectoderm clusters into two. **b**, Clustree flowchart by resolutions 0.1 to 0.4 in rows showing how higher resolutions generate more clusters. **c**, Prior to selecting clusters for merging in IS006 (WA09). **d**, The merging of clusters three and four from IS006 (WA09). After providing a username and/or a comment, the user can select the “Save to database”, which will automatically store the updated metadata in the RDS to be used for updated analysis.

**Extended Data Fig. 8: scRNA tab used to group cells by gene expression**

The mean of two genes per cell were selected here (SOX17 and HES4) and overlaid onto a UMAP. The user can either lasso cells to include them in a given set or define the minimum and maximum of gene expression for the targeted genes to select those cells for a given set. After providing a username and/or description of the type of grouping, the user selects “Save to database” to automatically store the updated metadata in the RDS to be used for updated analysis.

**Extended Data Fig. 9: iPSC Profiler module scores**

**a**, Endoderm module scores are highest in the endoderm cluster. **b-c**, the same effects for the ectoderm and mesoderm clusters with their respective modules. **d**, Housekeeping module with approximately even scores across all clusters. **e**, Heatmap module scores from both ScoreCard and iPSC Profiler indicating similar scores for pluripotency, ectoderm, mesoderm, and endoderm.

## Methods

The primary workflow is as follows: loading data from the existing bulk database or custom upload, Data structure, Analysis and the iPSC Profiler (Fig. 1). Extending from two previous frameworks, IRIS-EDA and scClustViz, there are many new feature enhancements for analysis that can be finely adjusted by the user in a highly interactive fashion^12, 18^.

We included two main additional features for bulk RNA-Seq: removing outliers and batch correction. Outliers can be identified by selecting “Subset data” then “Identify outliers” which creates a PCA. The user can select specific samples or technical replicates to remove. If bulk samples from different sequencing libraries are selected, batch correction is run by default using the RUVSeq RUVg algorithm with an empirical set of housekeeping genes^19^. We chose RUVg batch correction because it does not dampen biological signals as aggressively as other methods, such as ComBat-seq^20^. RUVg does not require a balanced experimental design with the same samples in each batch, making it highly convenient for merging disparate sequencing libraries. If batch correction was performed, the weighting variable is included first in the design as recommended by Risso *et al*.^19^.

We included five main features for single-cell data: 1. calculate multiple nearest-neighbor resolutions to find ideal cluster assignment, 2. improved visualizations using clustree for resolution selection, with converted scClustViz plots to ggplot2 and lasso selection for Seurat-based plots, 3. GSE, 4. options to manually combine nearest-neighbor clusters and create updated clusters based on the expression of selected gene(s), and 5. the iPSC Profiler tool. For both bulk and scRNA data, the user can merge samples across multiple experiments in the database or their own uploaded data, allowing comparisons to previously published data. For bulk data we have also included batch correction, which allows for the rapid comparison across datasets with reduced technical confounding factors. To exemplify the utility of SEQUIN, we describe bulk and single-cell analysis of human embryonic stem cells (WA09, WiCell, Madison, WI) and iPSCs (LiPSC-GR1.1, NIH Common Fund) that were differentiated into ectoderm, endoderm, and mesoderm using standardized kits (STEMCELL Technologies) as previously described^7^.

As the number of cells increases for scRNA experiments, so do the limitations within the R environment. While it is possible to load upwards of 100,000 cells into R, this will cause problems in R/Shiny. While we can confirm that we are able to load and analyze 55,074 cells in SEQUIN, it comes with significant wait time for loading plots, tables, and complete analyses. As scRNA datasets become even more prevalent and larger, this will also be a limitation of the app.

Although we included the option to batch correct bulk RNA-seq datasets, we did not include batch correction for scRNA datasets. Differential gene expression tests are sensitive to batch effects, but most approaches can only control for simple batch structure^21^. More complex experimental design such as unbalanced samples or uneven total cell counts across datasets can be problematic. We tested several batch corrections approaches on various scRNA datasets as well as simulated data, and we concluded that the majority of these methods completely dampened true biological signal rather than simply removing batch effects.

The iPSC Profiler was developed as a tool to quickly visualize the expression of gene sets in samples or cells or the score for a given gene module. The score is directly computed from Seurat’s AddModuleScore function, developed by Tirosh *et al*. 2016^22^. Since our data focuses on stem cells and iPSCs, genes from the ScoreCard were included in addition to longer literature and internal experimental based gene lists for the three primary tissue lineages ^16^. The user may choose either the canonical or expanded modules for fast assessment of early lineage attainment, as indicated by module name. For a brief comparison of the module score differences for trilineage, see Extended Data Fig. 9e.

## Data availability

Six publicly available experimental RNA-seq datasets are included in the app for training and demonstration purposes. The datasets generated by the Stem Cell Translation Laboratory (SCTL) were derived from hESCs (WA09) and iPSCs (LiPSC-GR1.1): “ISB003” (WA09 differentiated into three germ layers); bulk RNA-seq data “ISB008” (WA09 and LiPSC-GR1.1) from ref. 23; and four scRNA datasets from ref. 7: “IS020” (WA09 cultured manually and robotically); “IS018” (WA09 differentiated into neurons), “IS006_WA09” (WA09 differentiated into three germ layers); “IS0006_11” (LiPSC-GR1.1 differentiated into germ layers). A small RNA-Seq dataset “example_bulk” is from WI-38 fibroblasts with and without TGF-β treatment on days 1 and 20^24^. This dataset was selected because it is from human-derived samples, has a simple and balanced experimental design, and includes sufficient metadata to demonstrate all features and statistical models available for bulk RNA-seq in SEQUIN. A small scRNA-seq dataset “example_sc” was obtained from the IRIS-EDA GitHub repository (https://github.com/OSU-BMBL/IRIS) and consists of human preimplantation embryos and hESCs at different passages^18^. This dataset was selected because it is from human-derived samples and is small but sufficient for quickly demonstrating all features available for scRNA-seq in SEQUIN.

## Code availability

A publicly hosted version with example datasets from SCTL and the literature is available at https://sequin.ncats.io/app/. All code and a stand-alone package that can run locally is available here: https://github.com/ncats/public_sequin.

