## Supplementary figures and images for "SEQUIN: rapid and reproducible analysis of RNA-seq data in R/Shiny"

### Ext Data Fig 1

#### Extended Data Figure 1 (Weber et al.)

a

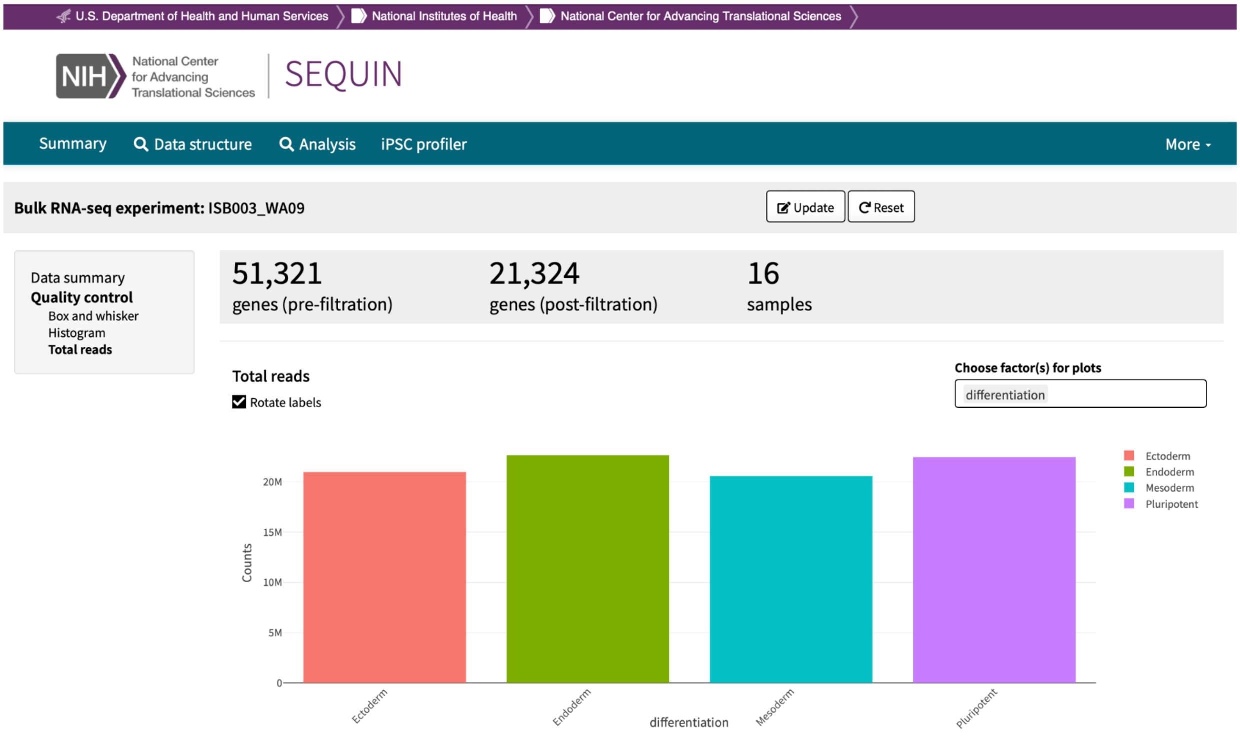


b

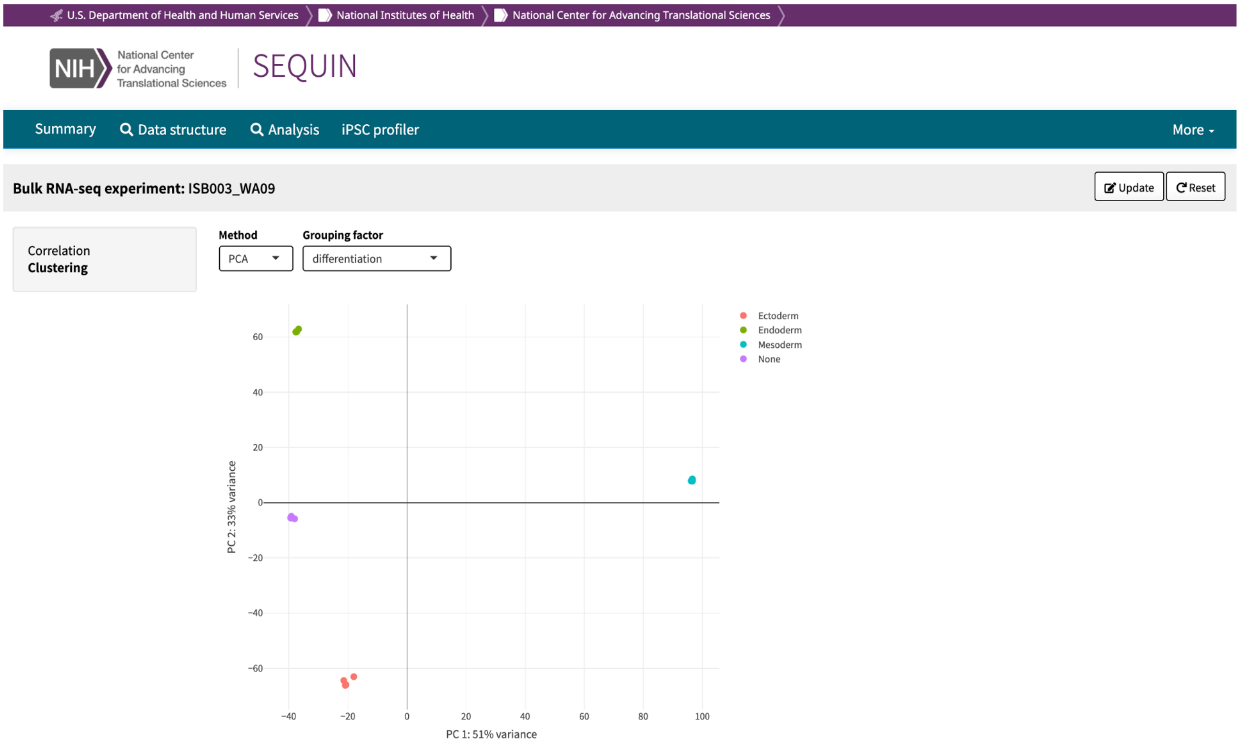

### Ext Data Fig 2

#### Extended Data Figure 2 (Weber et al.)

**a**

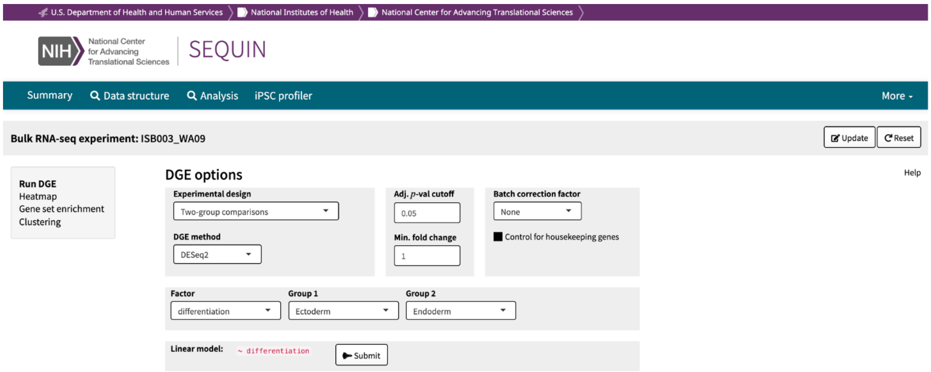


**b**

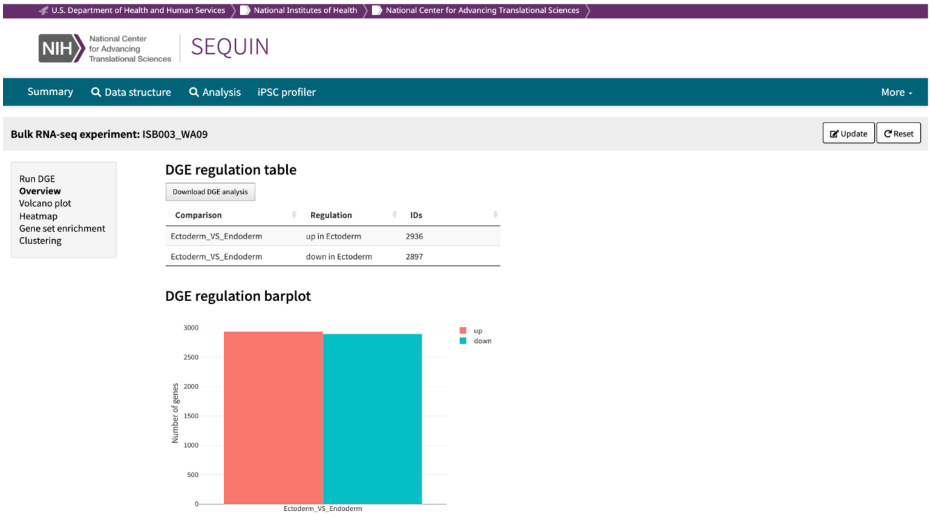


**c**

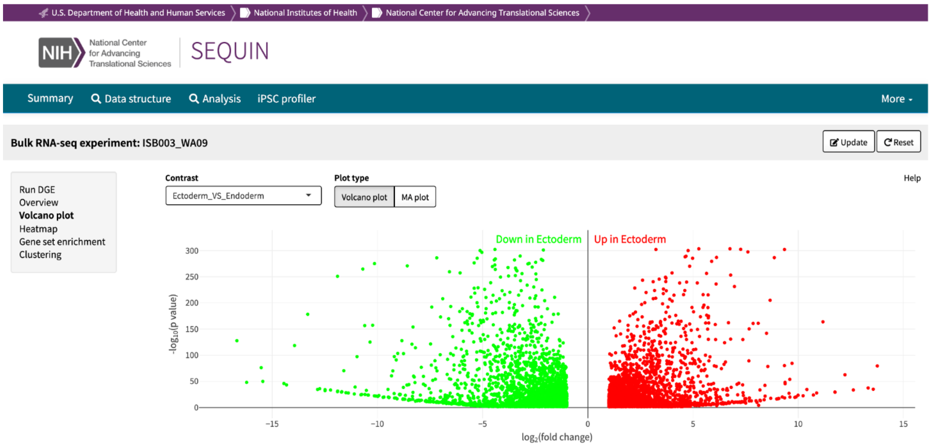

### Ext Data Fig 3

#### Extended Data Figure 3 (Weber et al.)

**a**

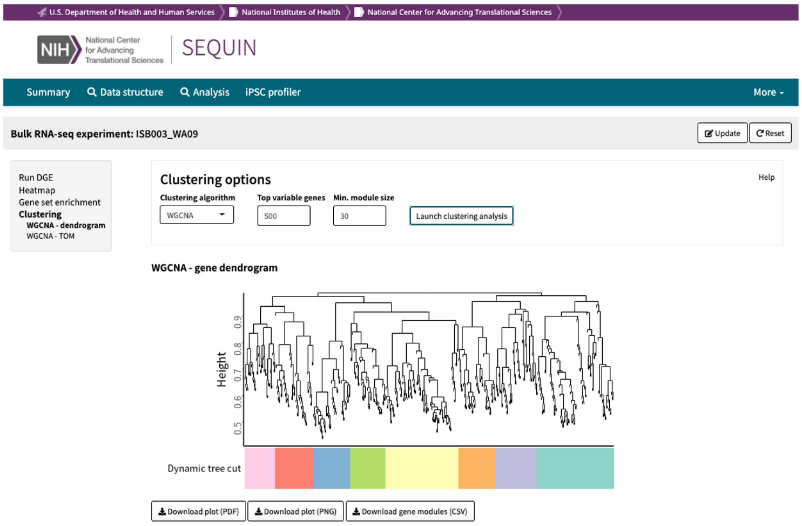


**b**

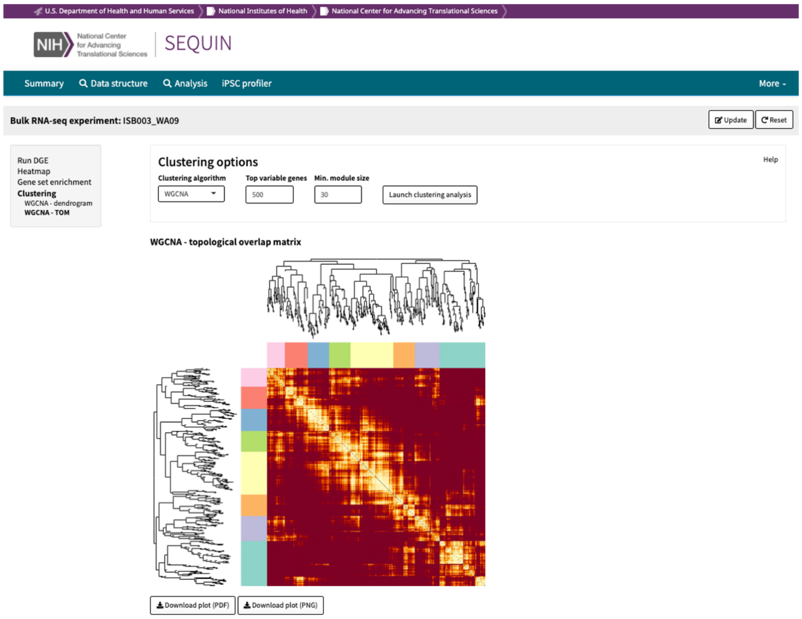


**c**

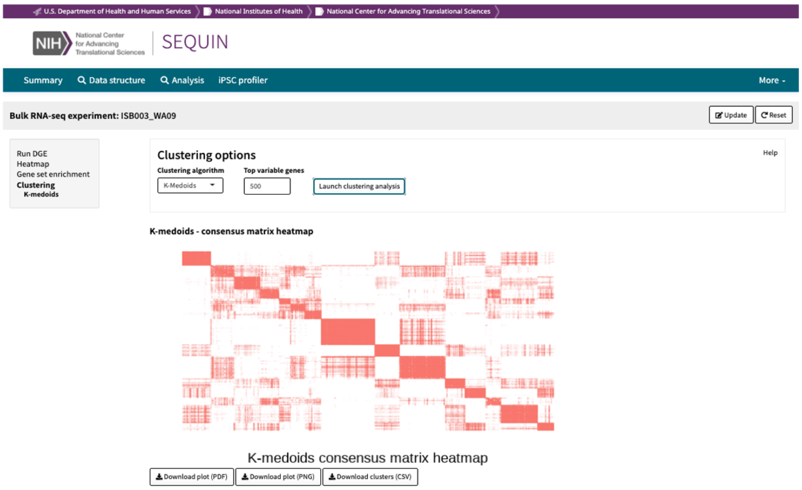

### Ext Data Fig 4

#### Extended Data Figure 4 (Weber et al.)

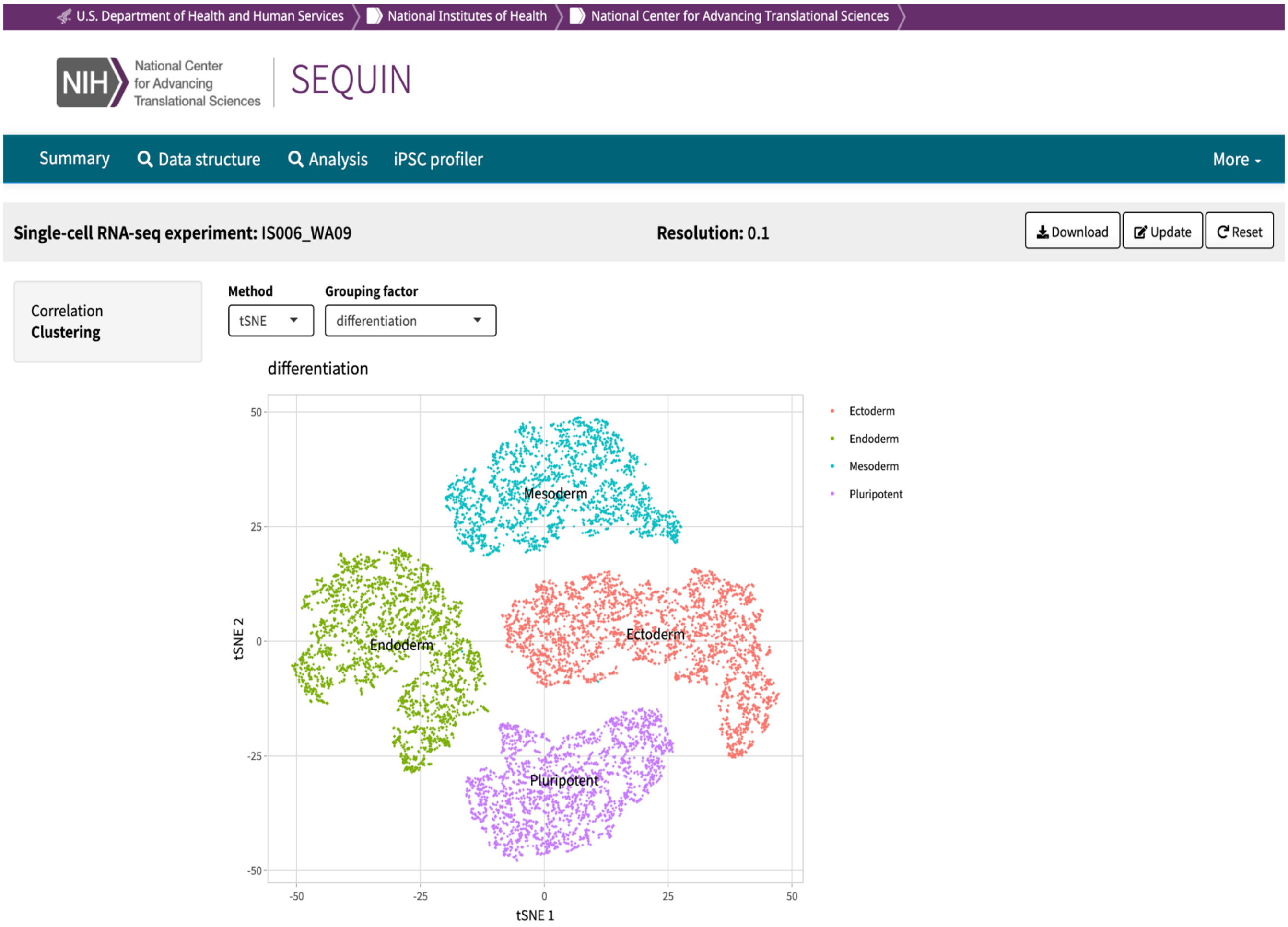

### Ext Data Fig 5

#### Extended Data Figure 5 (Weber et al.)

**a**

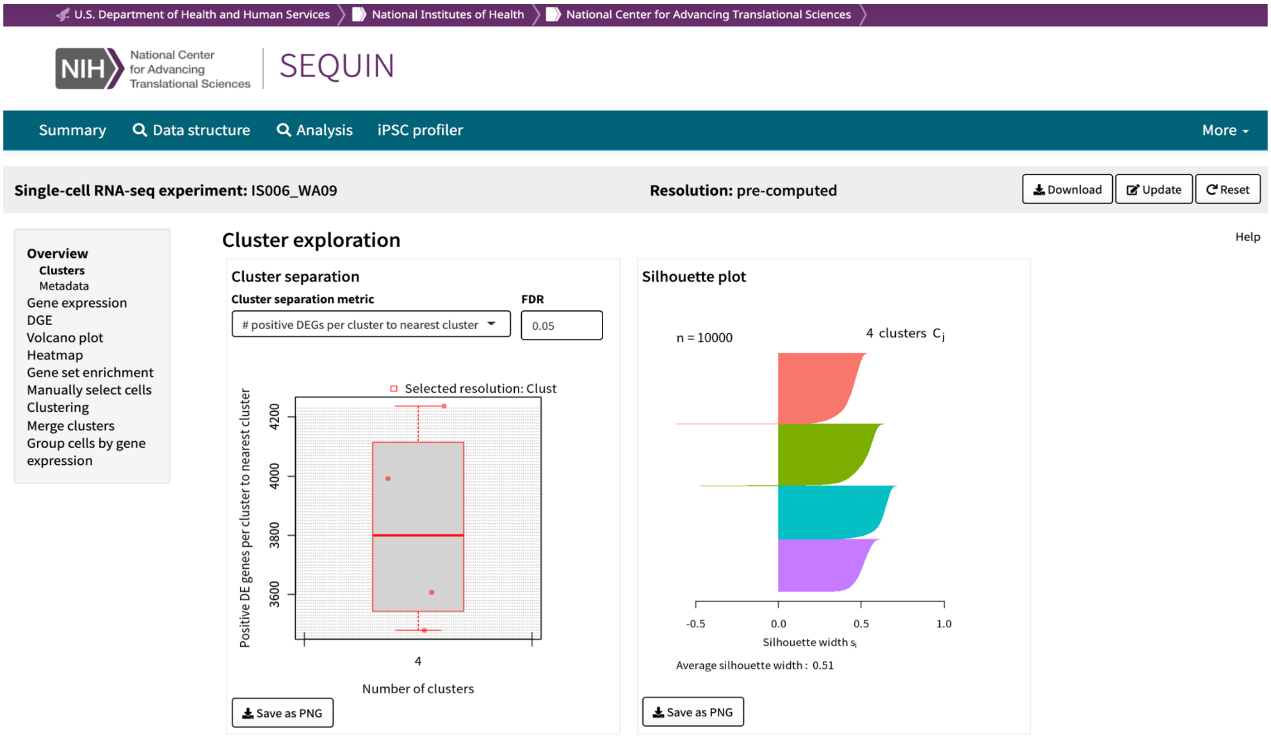


**b**

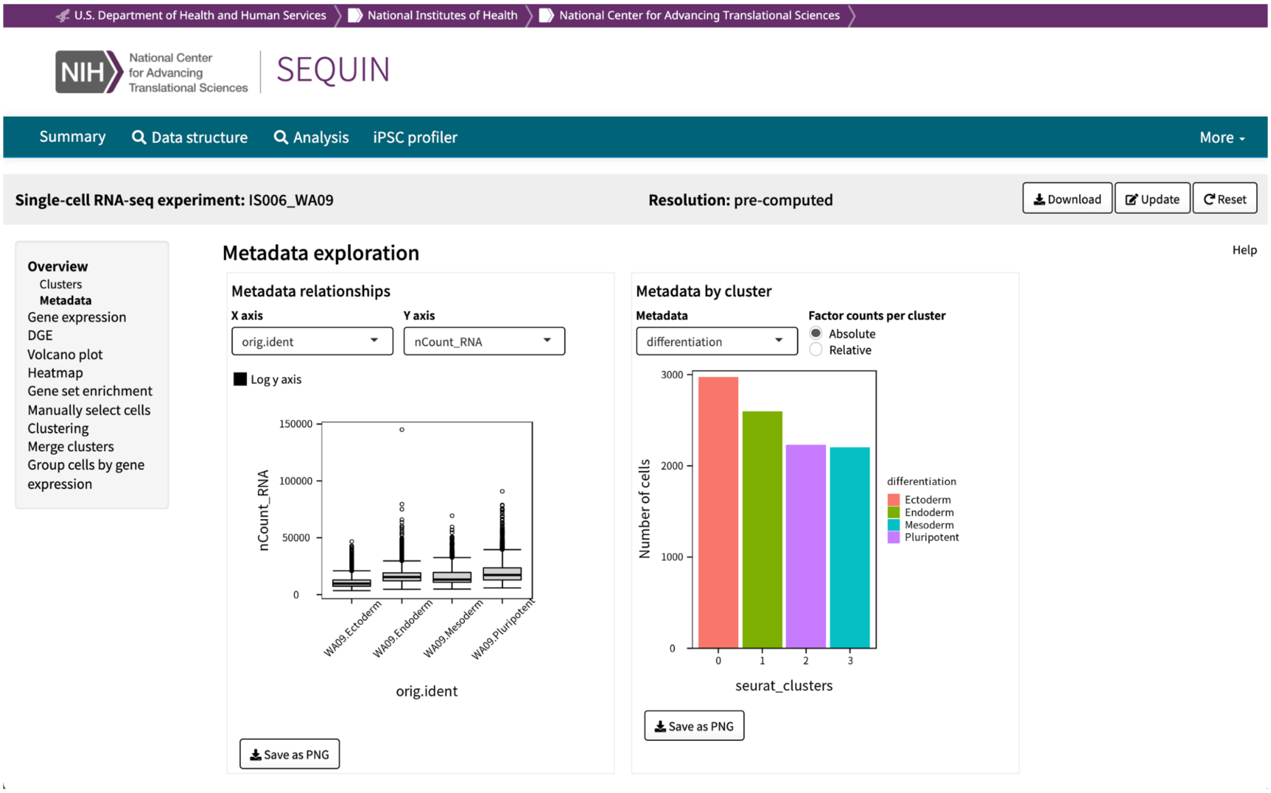

### Ext Data Fig 6

#### Extended Data Figure 6 (Weber et al.)

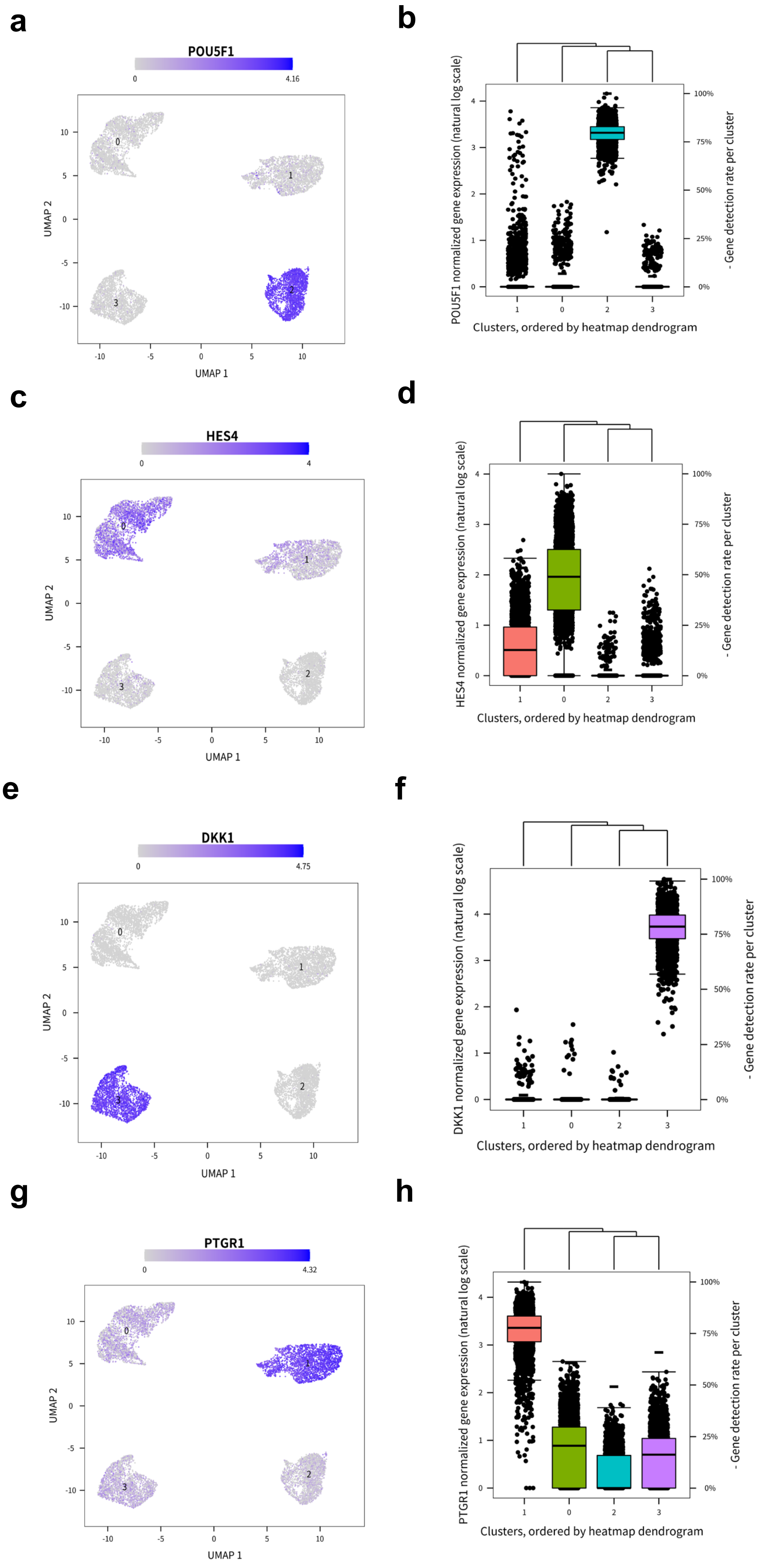

### Ext Data Fig 7

#### Extended Data Figure 7 (Weber et al.)

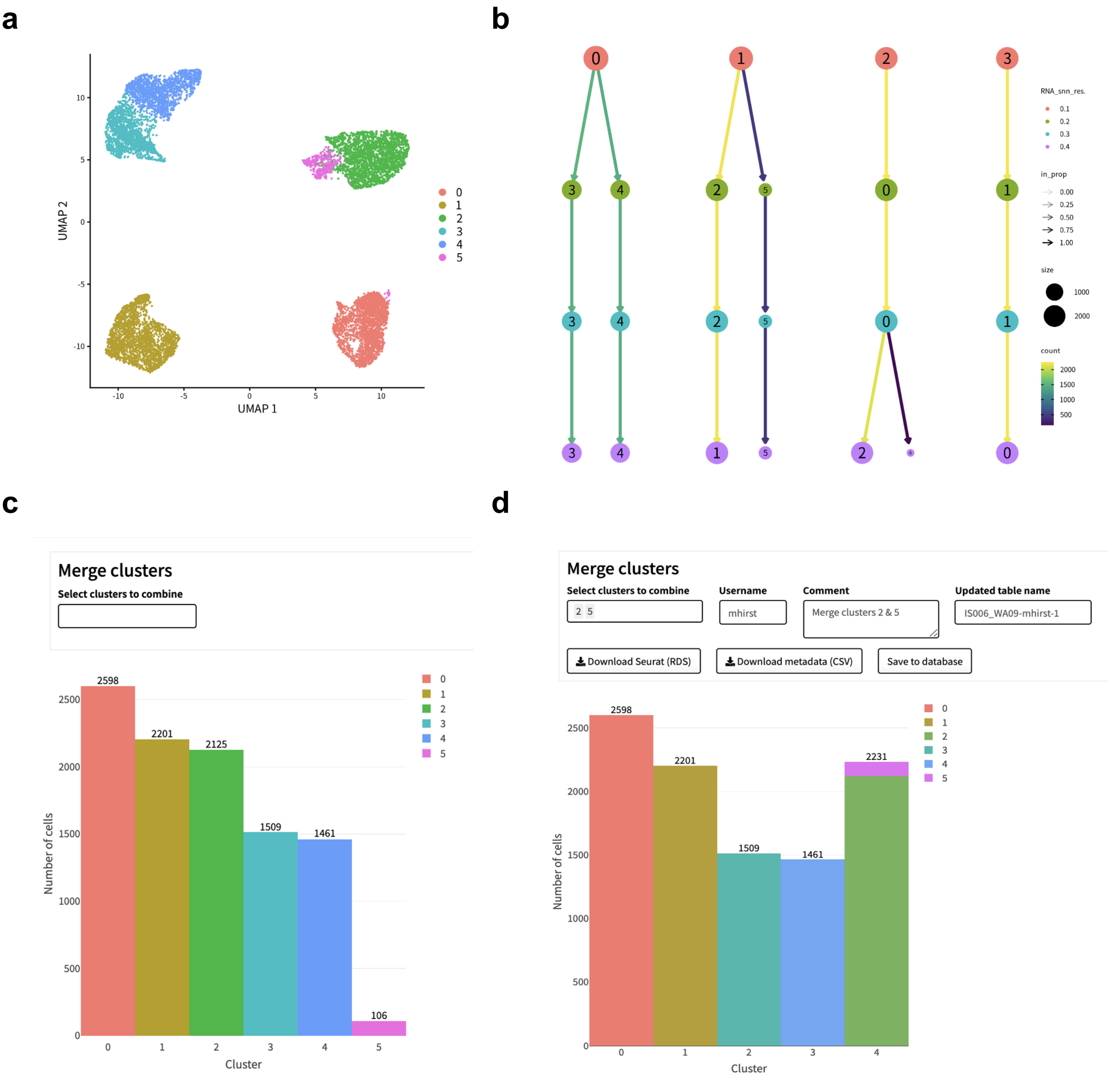

### Ext Data Fig 8

#### Extended Data Figure 8 (Weber et al.)

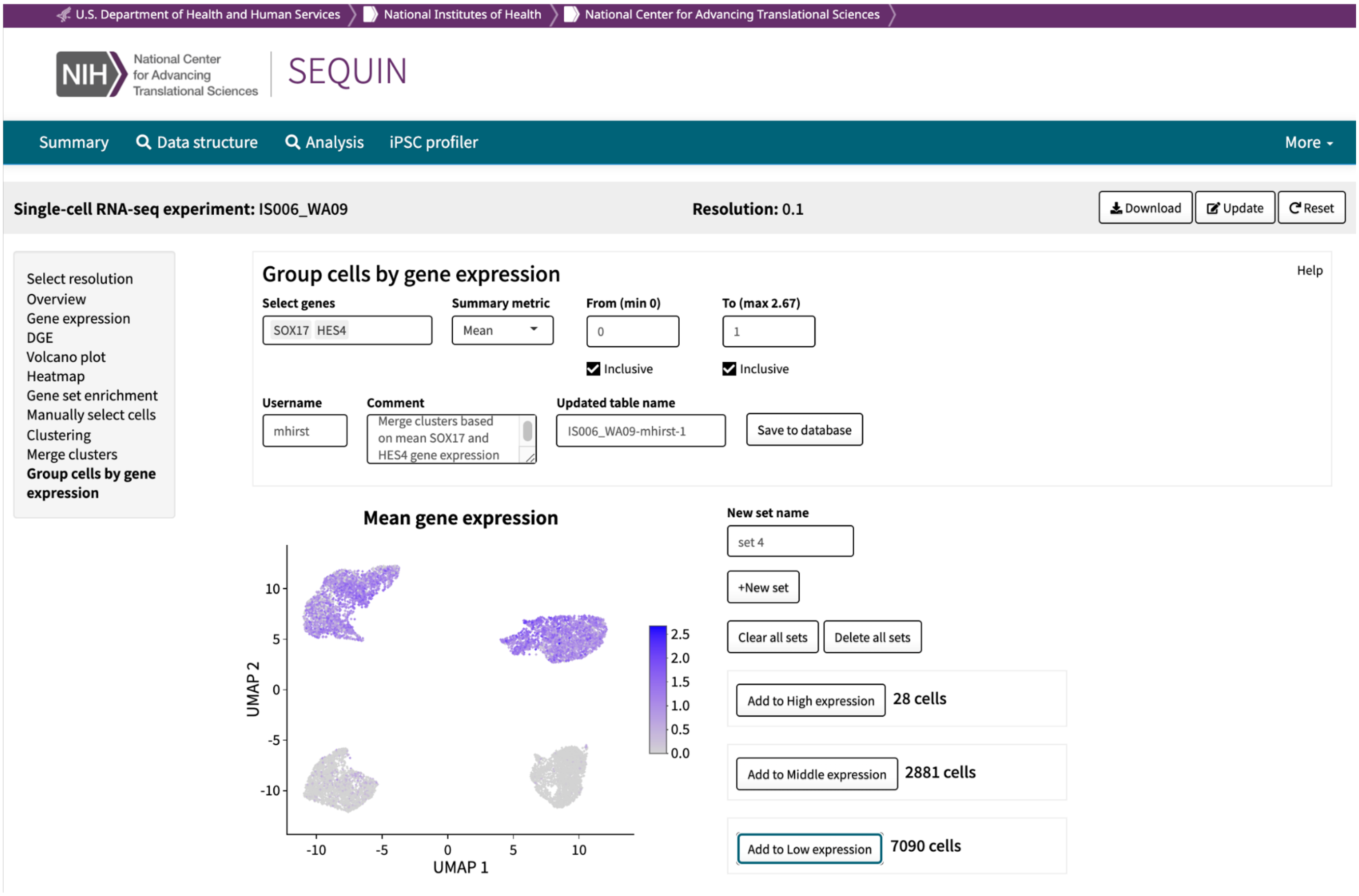

### Ext Data Fig 9

#### Extended Data Figure 9 (Weber et al.)

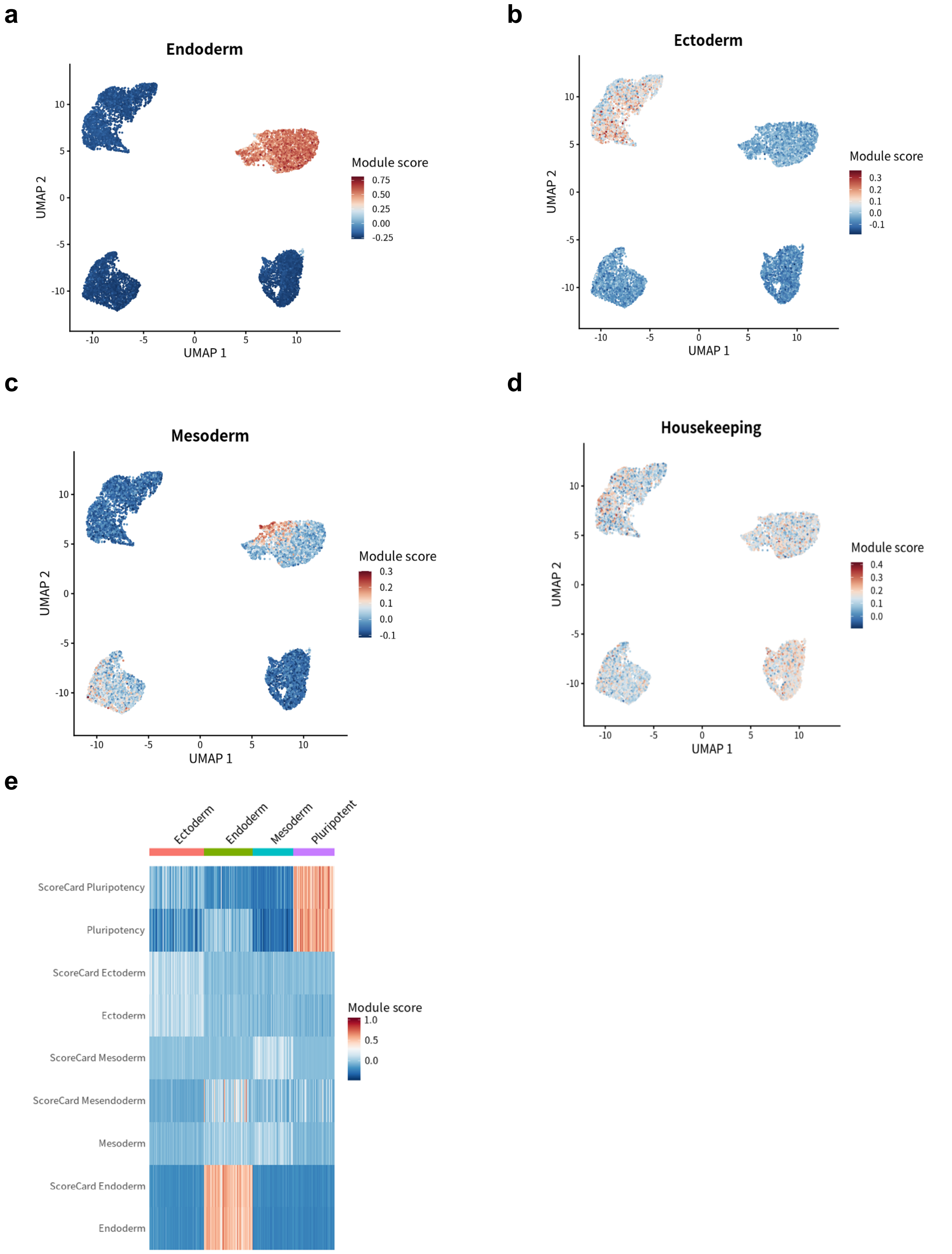
